# Creating a neuroprosthesis for active tactile exploration of textures

**DOI:** 10.1101/594994

**Authors:** Joseph E. O’Doherty, Solaiman Shokur, Leonel E. Medina, Mikhail A. Lebedev, Miguel A. L. Nicolelis

**Author notes:** JEO and SS have contributed equally to this work. Corresponding author: Miguel A. Nicolelis, 210 Research Drive, Box 103905 Dept of Neurobiology, Duke University, Durham, NC 27710.

## Abstract

Intracortical microstimulation (ICMS) of the primary somatosensory cortex (S1) can produce percepts that mimic somatic sensation and thus has potential as an approach to sensorize prosthetic limbs. However, it is not known whether ICMS could recreate active texture exploration—the ability to infer information about object texture by using one’s fingertips to scan a surface. Here we show that ICMS of S1 can convey information about the spatial frequencies of invisible virtual gratings through a process of active tactile exploration. Two rhesus monkeys scanned pairs of visually identical screen objects with the fingertip of a hand avatar, controlled via a joystick and later via a brain-machine interface, to find the one with denser virtual gratings. The gratings consisted of evenly spaced ridges that were signaled through ICMS pulses generated when the avatar’s fingertip crossed each ridge. The monkeys learned to interpret these ICMS patterns evoked by the interplay of their voluntary movements and the virtual textures of each object. Discrimination accuracy across a range of grating densities followed Weber’s law of just-noticeable differences (JND), a finding that matches normal cutaneous sensation. Moreover, one monkey developed an active scanning strategy where avatar velocity was integrated with the ICMS pulses to interpret the texture information. We propose that this approach could equip upper-limb neuroprostheses with direct access to texture features acquired during active exploration of natural objects.

## Introduction

Sensory neuroprostheses offer the promise of restoring perceptual function to people with impaired sensation ^1,2^. In such devices, diminished sensory modalities (e.g., hearing ^3^, vision ^4,5^, or cutaneous touch ^6–8^) are reenacted through streams of artificial input to the nervous system, typically using electrical stimulation of nerve fibers in the periphery or neurons in the central nervous system. Restored cutaneous touch, in particular, would be of great benefit for the users of upper-limb prostheses, who place a high priority on the ability to perform functions without the necessity to constantly engage visual attention ^9^. This could be achieved through the addition of artificial somatosensory channels to the prosthetic device ^1^. Such an approach would endow persons suffering from limb loss ^10–12^, paralysis ^1,13^ or somatosensory deficits with the ability to perform active tactile exploration of their physical environment and aid in dexterous object manipulation ^14–17^.

Previously we demonstrated that motor and sensory functions could be simultaneously enacted though a bidirectional neuroprosthetic system, called a brain-machine-brain interface (BMBI)^18^. In that demonstration, the active exploration enabled by our BMBI-driven neuroprosthesis used a limited and fixed set of ICMS temporal patterns to generate artificial sensory inputs that mimicked the sense of flutter-vibration. However, it remained unclear whether the same approach could generalize to allow the use of natural haptic exploratory procedures, where a person identifies the texture of objects and materials by scanning them with the fingertips.

Normal haptic exploration of objects involves several stereotypic procedures, such as static contact for temperature sensation, holding for weight, enclosure for gross shape, pressure for hardness, contour following for exact shape and lateral fingertip motion for texture ^19^. Here we developed a neuroprosthetic paradigm for restoring the sensation of fingertip motion against texture. We hypothesized that ICMS pulses generated by exploratory movements over virtual gratings and delivered to primary somatosensory cortex (S1) would allow discrimination of texture coarseness.

## Results

### Active texture encoding

Two rhesus monkeys (monkey M and monkey N) were chronically implanted with multielectrode cortical arrays ^18^ (Supplementary Fig. S1). These animals explored virtual objects on a computer screen using a realistic upper-limb avatar (Supplementary Fig. S2), which they operated manually with a joystick (Fig.1a) or using a BMI. On each trial, a pair of rectangles appeared either on the left or on the right side of the screen. The rectangles were visually identical, but each was associated with an invisible tactile grating whose properties were signaled by charge-balanced ICMS pulses applied to S1 (a region exhibiting left forearm receptive fields for monkey M and left lower-limb receptive fields for monkey N). Each grating consisted of evenly spaced vertical ridges, which were invisible to the monkeys. The spatial frequency of the ridges, *f*, ranged from 0.5 to 4.0 ridges/cm; an untextured object with no ridges (*f* = 0 ridges/cm) was also presented on some trials.

**Fig. 1.**
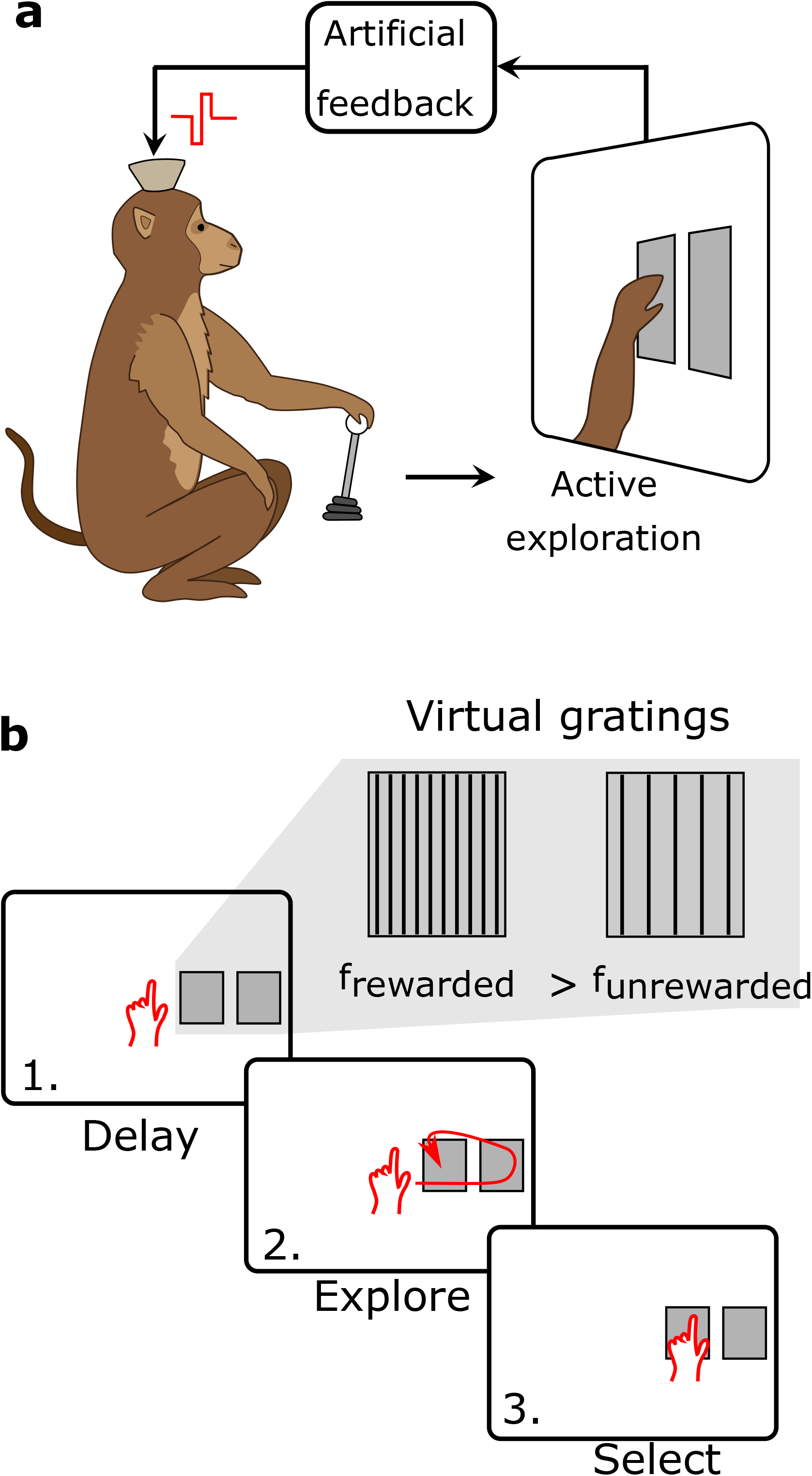
The artificial texture paradigm. (**a**) A monkey is seated before a display on which an avatar arm and two identical objects are projected. Artificial tactile feedback about the virtual gratings associated with each object is delivered to populations of S1 neurons via temporal patterns of ICMS as the monkey actively scans each object. (**b**) Trials commenced with a random delay interval (1) when the monkey held the index finger of the avatar in the center of the screen. Next, was the exploration interval (2). Two rectangular objects appeared, and the monkey scanned these objects with the index finger of the avatar hand. Each object had an associated virtual grating of vertical lines, which were invisible to the monkey. A pulse of ICMS was delivered to a pair of electrodes in S1 with each crossing of the avatar index finger over a line in one of the gratings. The trial was completed when the monkey indicated its selection (3) by holding the avatar hand over one of the objects for a hold interval. The reward was delivered if the monkey selected the object with the higher virtual grating frequency (inset); selecting the object with the lower grating frequency ended the trial without reward.

The behavioral task required the monkeys to probe the rectangles with the avatar’s fingertip, determine which of the two had a higher *f*, and to hold the avatar over that object for the required interval, 2 s in most cases (Fig. 1b). The artificial sensation was encoded by delivering a charge-balanced ICMS pulse each time the avatar fingertip crossed a ridge in a grating. Thus, the pulse-trains of ICMS delivered on any given trial provided an artificial signal that depended on the interplay between the movements of the avatar and the *f* of the textures of the explored objects (Supplementary Movie S1). Movements at a constant velocity across a grating with a given *f* produced an ICMS pulse train with a constant temporal pulse rate (Fig. 2a). Movements at a faster velocity across the same grating produced a pulse train with a correspondingly higher pulse rate (Fig. 2b). Irregular movements produced temporally varying ICMS pulse trains (Fig. 2c). The objects’ adjacent spacing on the screen encouraged the monkeys to rapidly shift the avatar from one object to the other and determine which one had a denser grating. The monkeys were permitted to explore the objects in any sequence and enter each object multiple times, to accumulate evidence, before making the selection. Accordingly, the monkeys could select an object on the first pass (Fig. 2d,e) or employ several explorations of individual objects (Fig. 2f) before making a final selection. Prior to these experiments, these monkeys participated in other studies ^18,20,21^ and became proficient in using the joystick and the hand avatar and making decisions using ICMS pulse trains. However, none of the previous experiments employed the particular ICMS encoding rule or the texture scanning paradigm presented in the current study.

**Fig. 2.**
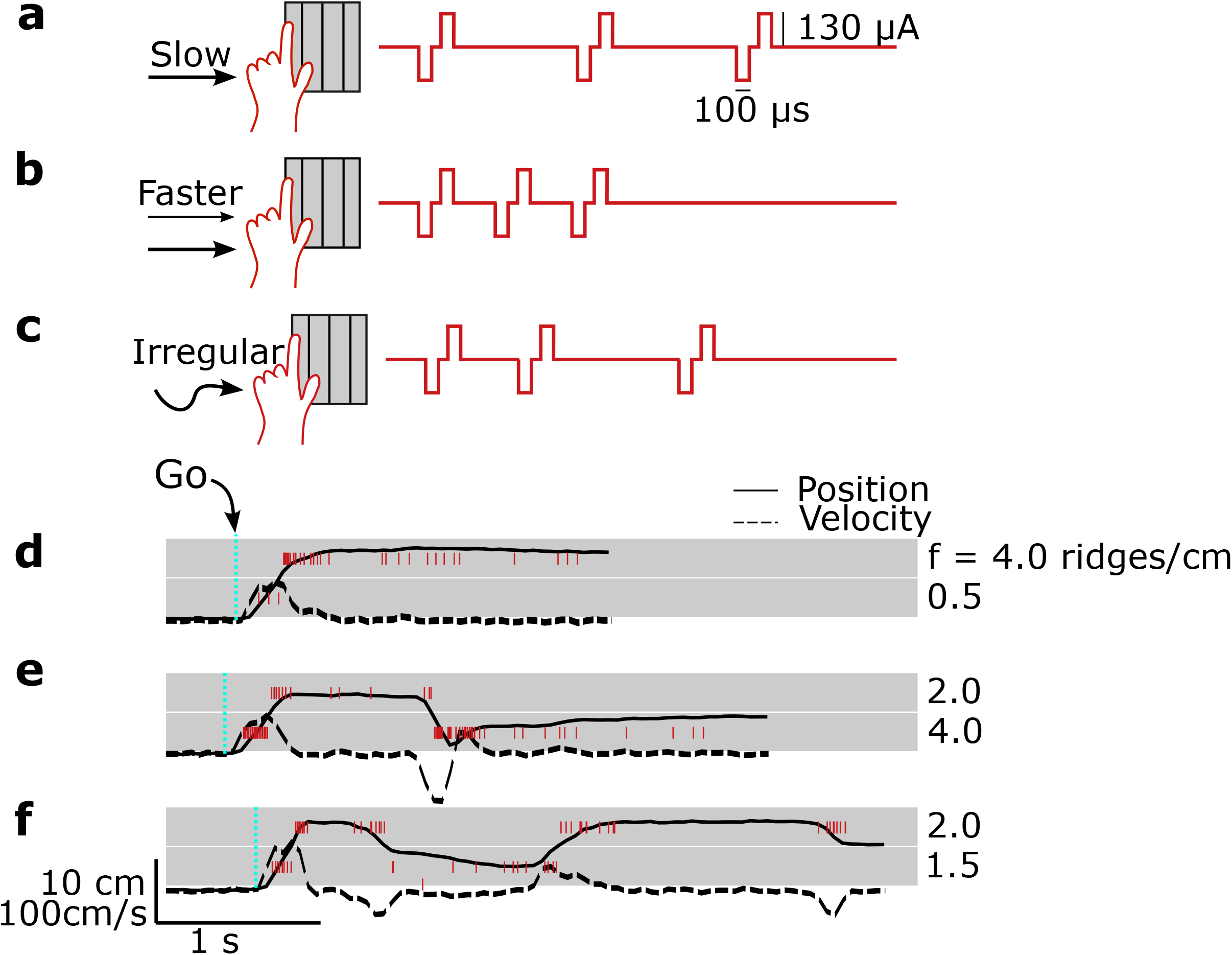
The precise temporal pattern of ICMS delivered on any trial depended both on the intrinsic spatial frequency of each object’s virtual grating as well as the velocity with which the monkey scanned each object. For a grating with a given spatial frequency, slow scanning (**a**) would produce a lower ICMS pulse rate than faster scanning (**b**). Irregular scanning (**c**) of a grating produced irregular ICMS pulse trains. All other features of the pulse train (e.g., current amplitude and pulse width) were fixed. (**d-f**). Examples of trials for three values of *Δf*: (**d**) 3.5 (4.0 vs 0.5) ridges/cm, (**e**) 2.0 (2.0 vs 4.0) ridges/cm, and (**f**) 0.5 (2.0 vs 1.5) ridges/cm, respectively. Traces indicate the x-component of the avatar position (solid lines) and velocity (dashed lines). Gray rectangles indicate the position and horizontal dimension of the objects. Red vertical lines indicate single pulses of ICMS. Trials started with a randomized hold-time (200-2000 ms); a Go cue informed the monkey of the beginning of the exploration interval.

### Active texture discrimination

Both monkeys learned the task rapidly, reaching high-performance levels (71% of correct trials for monkey N, and 73% for monkey M) after 10 daily sessions of training (Fig. 3a,b). The average performance was above chance even in the first training session (64% for monkey N and 56% for monkey M). For monkey M, task difficulty was increased gradually, with a large difference in *f* introduced early in training, *Δf* ≥ 2 ridges/cm; *Δf* < 2 ridges/cm after 3 sessions and the full range from *f* = 0 to *f* = 3.5 ridges/cm and a minimum *Δf* = 0.5 by the end of the training. The range for monkey N was *f* = 0 to *f* = 3.5 ridges/cm at the onset of the training and *f* = 0 to *f* = 4 by the end. The minimum difference between textures, *Δf*, was maintained at 0.5 for all sessions. Figure 3 c,d shows the behavioral performance after learning (11 and 12 recording sessions for monkeys M and N, respectively). Both monkeys performed better on individual trials when presented with larger *Δf* between the two objects than for smaller *Δf*, as might be expected. However, we observed an additional scaling of discrimination difficulty that depended on the absolute scale of the spatial frequencies of the objects being compared. More specifically, the psychometric functions for both monkeys were steeper for larger values of Σ*f*, that is, steeper for the larger sum for the two objects being compared (Fig. 4a,b).

**Fig. 3.**
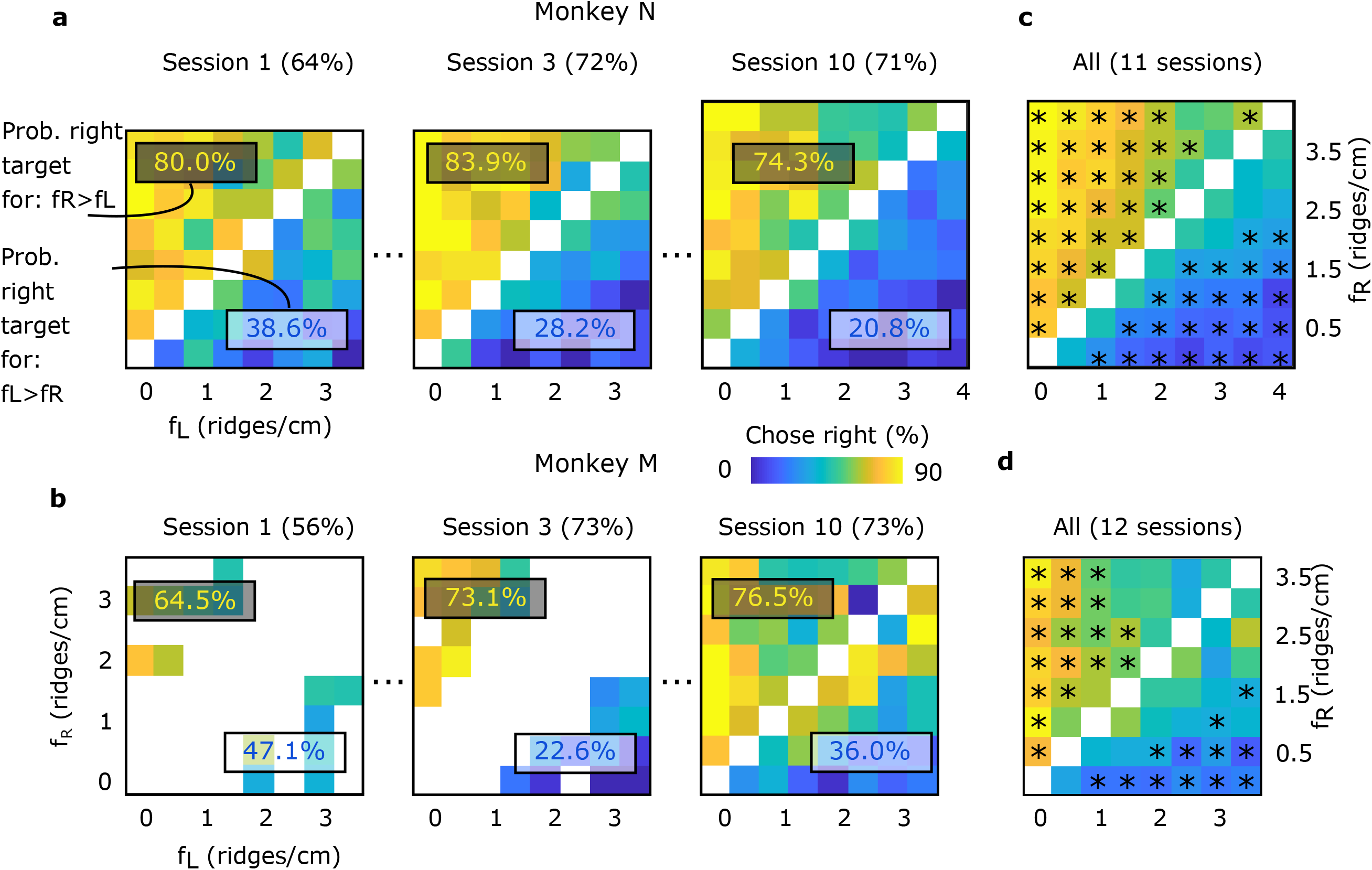
Monkeys discriminated spatial gratings based on self-generated temporal ICMS. (**a-d**) Percentage of trials for which the monkey chose the right-most object, parameterized by the spatial frequencies of the right and left objects, for both monkeys. (**a,b**) The success rate at the first session, and after three and after 10 sessions of training, is reported in parenthesis, and the average percentage of trails for which the right object was chosen when *f_R_>f_L_* and when *f_L_>f_R_*, reported in yellow and blue, respectively. (**c,d**) The performance for all sessions (11 sessions for monkey N, n=10412 and 12 for monkey M, n=5828); monkey M was not presented gratings with 4.0 ridges/cm. Asterisks indicate frequency-pair combinations which were discriminated significantly differently than chance (P<0.05, two-sided binomial test).

We quantified this phenomenon by estimating the just noticeable difference (JND), for each presented spatial frequency ^22^. We calculated, for each spatial frequency, the probability of choosing a second comparison frequency as a function of the unsigned delta between the standard stimulus and the comparison stimulus (Fig. S3). We found that the JND increased proportionally to *f* (Fig. 4c), consistent with the Weber–Fechner law ^23^ and Steven’s power law ^24^. The results for monkey M could be described by the linear function *JND*(*f*) = 0.47*f* + 1.06 (R^2^ = 0.63); *JND*(*f*) = 0.37*f* + 0.77 for monkey N (R^2^ = 0.95).

**Fig. 4.**
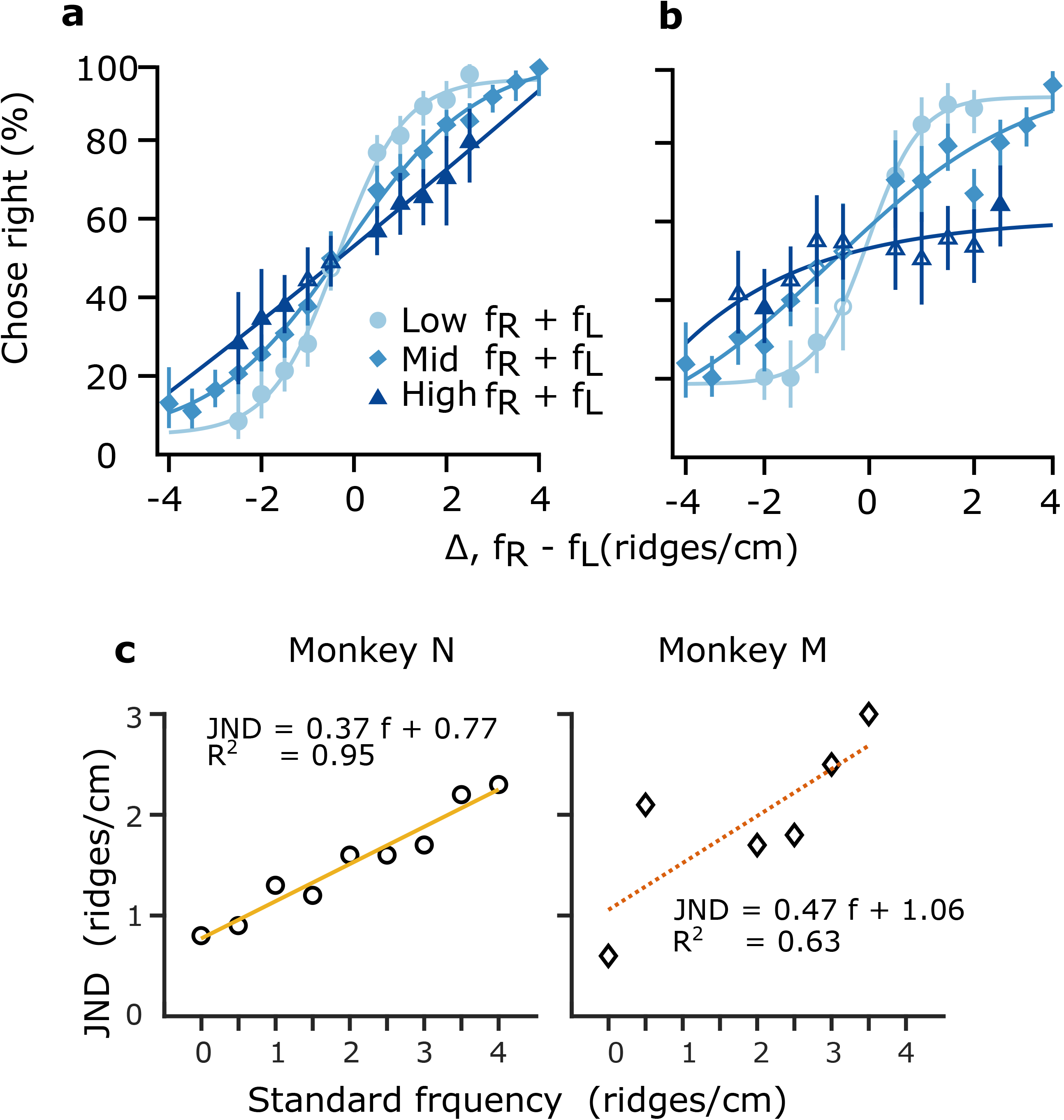
Psychometrics analysis of artificial texture discrimination. (**a,b**) Discrimination of spatial gratings obeys Weber’s scaling for (**a**) monkey N and (**b**) monkey M. Each point represents the percentage of trials for which the monkey chose the right-most object, parameterized by the difference in spatial frequencies for a pair of objects (*Δf, f_R_-f_L_*) and the sum of the spatial frequencies (*f_R_*+*f_L_*) for low (less than 2.5 ridges/cm, circles), mid (between 2.5 and 5 ridges/cm, diamonds) and high (greater than 5 ridges/cm, triangles) sums. Filled symbols indicate discrimination significantly different than chance (P<0.05, two-sided binomial test). Error-bars indicate 95% confidence intervals. Curves are the sigmoid lines of best fit. (**c**) Just noticeable differences (JNDs) for monkey M (diamonds) and N (circles), as a function of the standard frequency (detail of JND calculation for each standard frequency is shown on Figure S3; for monkey M JNDs for f =1 and f= 1.5 were undefined). Linear fits, the corresponding function and R^2^ for each graph.

There are a number of strategies that the monkeys could have used to compare the textures. One viable option would be to use a consistent velocity when exploring both objects so that any variation in ICMS pulse rate between the objects would be due to differences in spatial frequency alone. Further analysis revealed that this was not the case. Indeed, both monkeys used a distribution of speeds to sample the gratings (Fig. 5a) and could perform successful discriminations across the majority of their operating range (Fig. 5b)—only having difficulty when moving at very high speeds. Moreover, for the vast majority of trials, the average speeds used to scan the two objects differed, even within the same trial. Monkey M sampled the two objects with the same speed (delta speed < 1 cm/s) on fewer than 3% of trials, a finding that was not explained by the trial outcome (wrong trials: 2.41%, correct trials: 2.82%; Fig. S4). Monkey N used the same scanning speed for each target on only 3.85% of the trials (3.95% of the wrong trials, 3.81% of the correct trials).

**Fig. 5.**
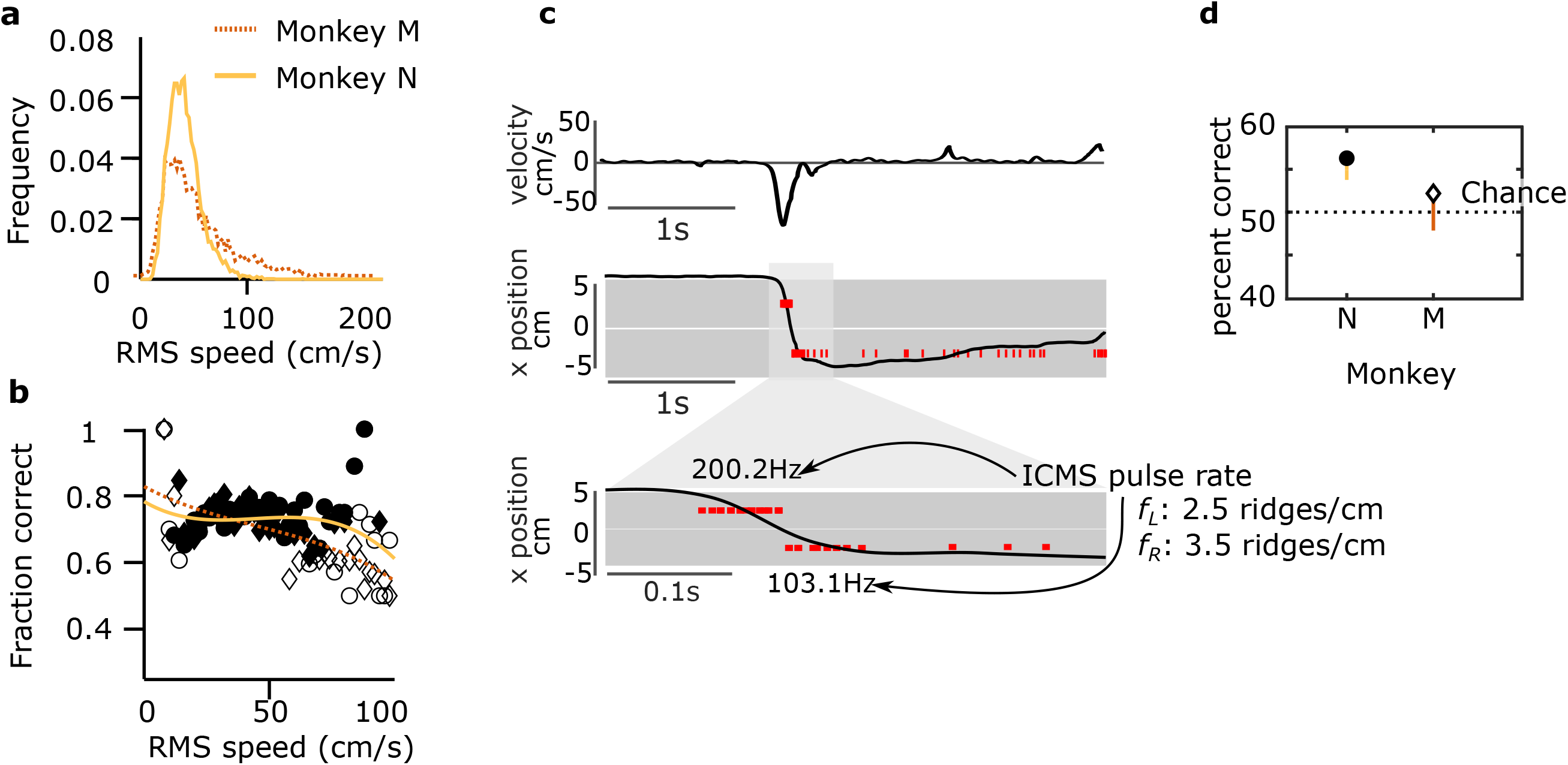
Texture perception and arm movement. (**a**) Distribution of per-trial RMS exploration speeds. (**b**) Percentage of trials performed correctly as a function per-trial RMS speed for monkey M and monkey N. Curves are 4^th^ order polynomial fits. Filled symbols indicate discrimination significantly different than chance (P<0.05, two-sided binomial test). (**c**) An example of a paradoxical trial with monkey N. First two graphs indicate the x-component of the avatar velocity and the x-position. Gray rectangles indicate the position and horizontal dimension of the objects; their corresponding spatial frequencies were 2.5 and 3.5 ridges/cm, respectively. Vertical red lines indicate single pulses of ICMS. ICMS pulse rate were calculated for bursts of stimulation (a burst of stimulation was considered when the velocity magnitude was maintained above 10 [cm/s]). (**d**) Success rate of the paradoxical trials. The chance level is reported with a black dashed line. Error-bars indicate 95% confidence intervals (one-sided binomial test).

This variability in arm movements was sufficiently large that, in some cases, the ordinality of spatial frequency of the textures was different from the ordinality of the ICMS pulses rates. An example of one of these apparently paradoxical trials is given in Figure 5c. For this trial, frequency of the right target (*F_R_* = 3.5 ridge/cm) was higher than the left (*F_L_* = 2.5 ridges/cm), but the actual ICMS pulse rate delivered for the left target was higher than for the right (left: 200.2 Hz versus right: 103.1 Hz). This occurred because a faster avatar speed was used to explore the left target as compared to the right. Despite this, the monkey was able to accurately choose the target with the higher spatial frequency in this example.

We found many of these apparently paradoxical trials (n=1231, 12% of all trials) for monkey N. The majority of these cases corresponded to frequency pairs with high FR+FL (Fig. S5). Monkey N’s success rate was significantly above chance for these trials (56.1%, P<0.001, one-tailed binomial test; Fig. 5d). There were fewer of these trials for monkey M (n=329, 6% of all trials). For these trials, monkey M’s performance did not reach significance (52.03%, P=0.25, one-tailed binomial test).

### Brain-machine-brain interface with active texture discrimination

Finally, we validated our stimulation paradigm in a closed-loop brain-machine-brain interface (BMBI) with monkey M. For this task, the monkey was allowed to move its arms, but the joystick was disconnected; instead the avatar arm—and task performance—was controlled via the decoding of 90 simultaneously recorded right-hemisphere M1 neurons (Fig. 6a). We found that monkey M was able to control the avatar arm to explore the objects with minimal movement of its physical hand as can be seen in the examples shown in Figure 6b. Moreover, when the hand did move, it made smaller movements with lower velocities than the simultaneous movements of the cursor during BMI trials (n=63 trials; Fig. 6c), but the monkey could still control the cursor using cortical activity alone (Supplementary Movie S2). The monkey retained the ability to accurately discriminate between the targets using the BMI; consistent with the non-BMI task, the monkey was significantly above chance in discriminating targets with Low Σ*f* (76%, P=0.02, one-sided binomial test), but did not reach significance for medium (65%, P=0.09) or high Σ*f* (40%, P = 0.21; Fig. 6d).

**Fig. 6.**
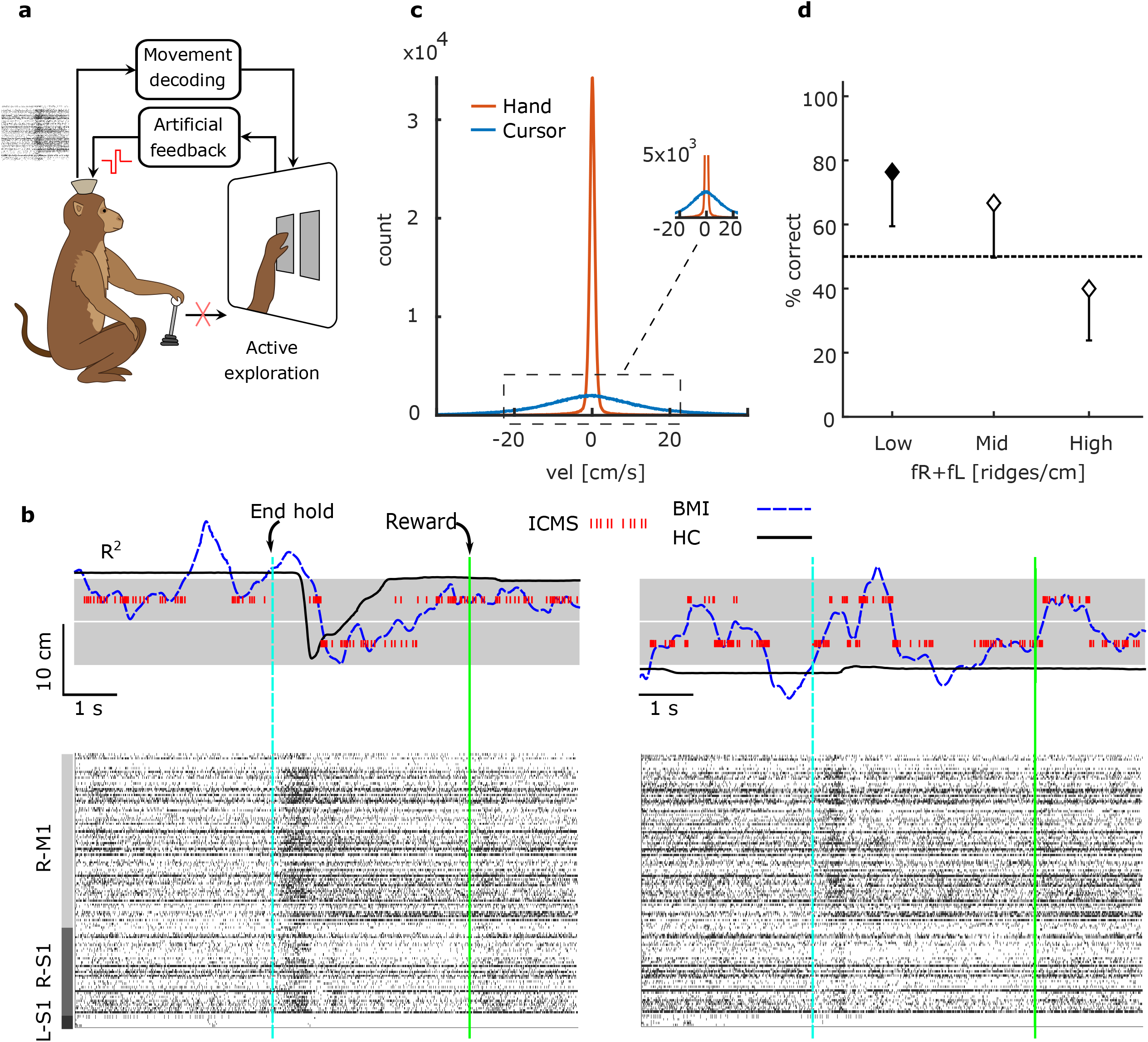
BMI results. (**a**) Same experimental paradigm as in Fig 1a, expect that the control of the avatar arm was done via decoding of motor intention from monkey’s motor cortex. The monkey has to hold the joystick during the task, was allowed to move the arm, but the joystick was disconnected (**b**) Examples of two BMI trials and corresponding raster plots. Blue dashed lines report the x projection of the brain-controlled cursor (BMI) and solid black line the monkey’s hand movement. ICMS pulses are shown with red vertical lines. Vertical dashed cyan line is the end of the hold time (or onset of exploration), and solid green line is the end of the trial and reward. Raster plots for the trials are grouped between 90 neurons in right hemisphere motor cortex area (R-M1), 47 neurons in right hemisphere sensory area (R-S1) and five neurons in the left hemisphere sensory area. (**c**) The distributions of velocity for the BMI controlled cursor (blue) distribution of the monkey’s hand movements (orange). The hand movement was measured via the joystick movement (using the same gain as for hand control trials) and the trial was aborted if the monkey released the joystick handle. (**d**) Percentage success for BMI controlled trials, parameterized by the sum of the spatial frequencies (*f_R_*+*f_L_*) for low (less than 2.5 ridges/cm), mid (between 2.5 and 5 ridges/cm) and high (greater than 5 ridges/cm) sums. Error-bars indicate 95% confidence intervals. Filled symbols are statistically different than chance (P<0.05, one-sided binomial test).

## Discussion

We have demonstrated a novel encoding strategy for texture representation using ICMS pulses in somatosensory cortex. Using this new approach, two animals were able to discriminate texture coarseness during active tactile exploration. Importantly, for this task, small variations of arm velocity changed the stimulation frequency; the interpretation of the texture, therefore, may have employed a dynamic integration of ICMS stimulation information with arm proprioception feedback or corollary discharge of motor and sensory cortical regions ^25^. The apparently paradoxical trials provided evidence for these possibilities: access to the movement command or proprioceptive feedback about the movement is necessary to disambiguate the exafference of the texture from the reafference due to movement.

We observed that both monkeys were better at discriminating textures when the overall spatial frequencies were small, consistent with the Weber-Fechner law ^26^, a phenomenon reported for numerous sensory modalities ^27^, including touch ^28^. Interestingly, this law was previously reported not to hold for the task of discriminating ICMS amplitude in primates ^29^ and humans ^13^. Our task, in contrast, required discriminating ICMS pulse rates, but, as it also used active exploration we cannot rule out the possibility that some aspect of the effect is due to the motor act itself.

Our tactile encoding scheme was effective for a single channel of independent tactile information—mimicking a single mechanoreceptor localized in the fingertip. This encoding scheme most closely resembles the rapidly adapting (RA) afferents of cutaneous somatic sensation ^30^: each pulse of ICMS was triggered by the intersection of the active zone of the avatar fingertip with a ridge on one of the gratings. However, there may be advantages of modeling a more slowly adapting type-1 (SA1) encoding on some additional channels. We believe that our encoding will be naturally extendable to arrays of mechanosensors embedded in the "skin” of a prosthetic limb, with each sensor connected to a channel of microstimulation in sensory cortex. For example, each feature in an object’s tactile microstructure could trigger a pulse-train of ICMS that persists for some finite duration. This type of encoding may allow an intuitive representation of the persistence of object-actuator contact interactions or complex representation of natural textures ^31^. However, a number of open questions remain, such as the optimal timescale or distribution of timescales for adaptation and whether the degree of adaptation must be matched to the properties of the specific neurons being stimulated. Work in primates ^6,32^ and rats ^33^ suggests that the plasticity of the brain will allow even a few channels of stimulation to become effective at providing a rich sensory experience, and complex spatiotemporal coding ^34^ with enough bandwidth to be clinically useful.

In our experiment, monkey N was superior to monkey M in perceiving small differences of texture coarseness. While it is possible that this difference was due to a better comprehension of the task by monkey N, it could also reflect the fact that the stimulation region for monkey N was in the leg area while for monkey M it was in the receptive fields of the same arm used to control the joystick. Therefore, it is possible that interference between feedback from natural somatosensory pathways (hand touching the joystick, proprioception) and S1 ICMS feedback made interpretation more difficult for monkey M. This indicates that further studies are necessary to determine, among other things, the best target in S1 for delivering ICMS that encodes tactile signals for future clinical neuroprosthesis. While delivering sensory feedback to an ethologically meaningful cortical area is likely important for the subject to assimilate any limb prosthesis as a natural appendage ^35–37^, the use of different somatosensory regions in the cortex may facilitate the sensory-motor integration and tactile acuity. Therefore, we suggest that it may be necessary to deliver artificial sensory feedback to multiple cortical regions simultaneously to achieve the best performance of such limb prostheses.

Recently demonstrated clinical neuroprostheses have used modulation of stimulation amplitude (or equivalently, pulse-width) to encode the perception of pressure, force or position ^8, 10, 38, 39^ Our approach is complementary—stimulation pulse timing encodes coarse texture—and could be combined with the amplitude encoding approach to convey multimodal percepts of pressure and texture. However some previous animal ^40^ and human ^41^ stimulation studies have provided indirect evidence that changes in pulse intensity (amplitude or pulse-width) may be perceptually indistinguishable from changes in pulse rate. Further experiments will be necessary to conclusively determine if this is the case or if there is in fact an extra degree of freedom that can be used to convey clinically relevant prosthetic sensations.

Finally, we demonstrated that our encoding strategy could be integrated within a closed-loop BMBI task. While the overall performance of the monkey for the BMBI task was lower than during arm-control, the monkey was still able to discriminate the artificial textures. This, along with the simplicity of our ICMS encoding, suggests that this approach could be used to equip clinical upper-limb neuroprostheses with direct access to the tactile features of the natural world.

## Online methods

All animal procedures were performed in accordance with the National Research Council’s Guide for the Care and Use of Laboratory Animals and were approved by the Duke University Institutional Animal Care and Use Committee.

### Subjects and Implants

Two adult rhesus macaque monkeys (*Macaca mulatta*) participated in the experiments (monkeys M and N). Each monkey was implanted with four 96-microwire arrays constructed of insulated stainless steel 304. Each hemisphere received two arrays: one in the upper-limb representation area and one in the lower-limb representation area of sensorimotor cortex. These arrays covered both M1 and S1; only microwires implanted in S1 were used for delivering ICMS in study. For the BMI task, we used recordings from the right hemisphere arm arrays as the monkey manipulated the joystick with the left arm. Within each array, microwires were grouped in two four-by-fours, uniformly spaced grids each consisting of 16 electrode triplets. The separation between electrode triplets was 1 mm. The electrodes in each triplet had three different lengths, increasing in 300-mm steps. The penetration depth of each triplet was adjusted with a miniature screw. After adjustments during the month following the implantation surgery, the depth of the triplets was fixed. The longest electrode in each triplet penetrated to a depth of 2 mm as measured from the cortical surface.

### Task

Each monkey sat in a primate chair, faced a computer screen and grasped a joystick with their left hand. The joystick handle contained an optical sensor to indicate when the monkey released it. The monkeys were trained to manipulate the joystick to control the movements of a left upper-limb primate avatar on the screen ^18,42^.

Each trial began with a circular target appearing in the center of the screen. The monkeys held an index finger of the avatar within this target for a random delay randomly drawn from a uniform distribution parameterized from 200 to 2000 ms. After this delay, the central target disappeared, and two rectangular object zones appeared on the screen. These appeared either both on the left side or both on the right side of the screen at a distance of 7 cm from the center. Both objects in the pair had the same width, (6 cm). The spacing between the objects was 0.1 cm.

Vertical square-wave gratings were superimposed on each of the objects. These gratings, which were not visible to the monkeys, were aligned on the center of each object and were parameterized by spatial frequency, *f*. When the index finger of the avatar crossed a single ridge in a grating, a pulse of ICMS was delivered to a pair of electrodes implanted in S1 cortex. In this way, the pattern of ICMS delivered depended on the velocity of the avatar and the intrinsic spatial frequency of each grating. The microstimulator was serviced at 100 Hz, which meant that for sufficiently fast velocities or high spatial frequencies, it could be possible that more than a single ridge was crossed in a 10 ms interval. If this occurred, we delivered N pulses at N*100 Hz, where N was the number of ridges crossed since the last clock cycle. This operation delivered the correct number of pulses at the correct rate, in expectation, at the cost of up to 10 ms of latency.

Symmetric, biphasic, charge-balanced, cathode-leading ICMS pulses were delivered in a bipolar fashion across pairs of microwires. The channels selected had clear sensory receptive fields in the left forearm (monkey M: two pairs of microwires) or left lower limb (monkey N: one pair of microwires). For monkey M, the cathodic and anodic phases of stimulation had a pulse width of 105 μs; for monkey N, the pulse phases were each 200 μs. The cathodic and anodic of the stimulation waveforms were separated by a 25 μs interphase interval. The pulse amplitudes were set to the minimal effective current, as found through psychometric measurements separately for each monkey ^43^.

Monkeys received a reward for selecting the object from the pair with the higher spatial frequency, *f*, drawn from:

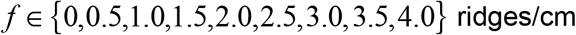

with the constraint that both objects did not share the same *f* on a single trial. Monkey M did not discriminate the gratings as reliably and so was not presented any gratings with the highest spatial frequency, 4.0 ridges/cm. The monkeys indicated their choice by holding the avatar over one of the objects for the hold interval (2 seconds for the hand control and 1 second for the BMBI task). Selecting the object with the higher *f* triggered the delivery of a fruit juice reward; selecting the object with lower *f* ended the trial without reward.

The objects could be explored in any sequence. Moreover, objects could be re-explored and re-compared multiple times in a trial. However, the avatar had to pass over both objects at least once per trial. Trials for which only a single object was explored were terminated without reward, even if the correct object was ultimately selected. Trials for which the monkey released the joystick handle at any time, selected the wrong object, made a selection without exploring both objects, or held the avatar outside of either of the objects for 10 s, resulted in the termination of a trial and penalty interval of 2 s for monkey M and 2.5 s for monkey N.

We employed correction trials. This meant that after an incorrect trial, the next one repeated with the same object locations and object-frequency identities. These correction trials were used to keep the monkeys motivated and to prevent them from acquiring systematic biases. As the rewarded object was known to the monkeys for correction trials, we excluded these trials from all analyses.

### BMI decoding

A 10th-order Unscented Kalman filter (UKF) was used for BMI predictions, using methods we previously described ^18,44^. The filter parameters were fit using the hand movements made while the task was performed using a joystick. The monkey was permitted to continue moving the joystick, but was only rewarded for target selections made with the brain-controlled cursor.

## Acknowledgments

We thank D. Dimitrov for conducting the animal surgeries, J. Fruh for the design of the monkey avatar, and G. Lehew, J. Meloy, T. Phillips, L. Oliveira, Amol Yadav and S. Halkiotis for technical support. This research was supported by DARPA N66001-06-C-2019, TATRC W81XWH-08-2-0119, and NIH Director’s Pioneer Award DP1OD006798 to MALN.

## Author contributions

JEO, MAL, and MALN designed the experiments. JEO, LEM, and MAL conducted the experiments. JEO, SS, MAL, and MALN analyzed data and wrote the paper.

## Materials & Correspondence

The custom code used for the experiments and the data that support the findings of this study is available from the corresponding author upon reasonable request. Requests should be addressed to

## References and Notes

1. Lebedev, M. A. & Nicolelis, M. A. L. Brain – machine interface□: past, present and future. 29, (2006).

2. Collinger, J. L., Gaunt, R. A. & Schwartz, A. B. Progress towards restoring upper limb movement and sensation through intracortical brain-computer interfaces. Curr. Opin. Biomed. Eng. (2018). doi:10.1016/j.cobme.2018.11.005

3. Wilson, B. S. et al. Better speech recognition with cochlear implants. Nature 352, 236–238 (1991).

4. Humayun, M. S. et al. Visual perception in a blind subject with a chronic microelectronic retinal prosthesis. Vision Res. 43, 2573–2581 (2003).

5. Normann, R. A., Maynard, E. M., Rousche, P. J. & Warren, D. J. A neural interface for a cortical vision prosthesis. Vision Res. 39, 2577–2587 (1999).

6. Fitzsimmons, N. A., Drake, W. & Hanson, T. L. Primate reaching cued by multichannel spatiotemporal cortical microstimulation. J. Neurosci. (2007). at <http://www.jneurosci.org/content/27/21/5593.short>

7. Romo, R., Hernández, A., Zainos, A. & Salinas, E. Somatosensorydiscrimination basedoncortical microstimulation. Nature 392, 387–390 (1998).

8. Tan, D. W. et al. A neural interface provides long-term stable natural touch perception. Sci. Transl. Med. (2014). doi:10.1126/scitranslmed.3008669

9. Atkins, D. J., Heard, D. C. Y. & Donovan, W. H. Epidemiologic Overview of Individuals with Upper-Limb Loss and Their Reported Research Priorities. J. Prosthetics Orthot. 8, 2 (1996).

10. Raspopovic, S. et al. Supplementary materials for: Restoring natural sensory feedback in real-time bidirectional hand prostheses. Sci. Transl. Med. 6, 222ra19 (2014).

11. Marasco, P. D., Schultz, A. E. & Kuiken, T. A. Sensory capacity of reinnervated skin after redirection of amputated upper limb nerves to the chest. Brain (2009). at <http://brain.oxfordjournals.org/content/132/6/1441.short>

12. Oddo, C. M. et al. Intraneural stimulation elicits discrimination of textural features by artificial fingertip in intact and amputee humans. Elife 5, 1–27 (2016).

13. Flesher, S. N. et al. Intracortical microstimulation of human somatosensory cortex. Sci. Transl. Med. 8, 361ra141–361ra141 (2016).

14. Moberg, E. Criticism and studvy of methods for examining sensibility in the hand*. Neurology (2012). doi:10.1212/wnl.12.1.8

15. Flanagan, J. R. & Wing, A. M. Modulation of grip force with load force during point-to-point arm movements. Exp. Brain Res. 95, 131–143 (1993).

16. Johansson, R. S. & Westling, G. Roles of glabrous skin receptors and sensorimotor memory in automatic control of precision grip when lifting rougher or more slippery objects. Exp. Brain Res. (1984). at <http://www.springerlink.com/index/M402455817334028.pdf>

17. Johansson, R. S. & Flanagan, J. R. Coding and use of tactile signals from the fingertips in object manipulation tasks. Nat. Rev. Neurosci. 10, 345–359 (2009).

18. O’Doherty, J. E. et al. Active tactile exploration using a brain–machine–brain interface. Nature 479, 228–231 (2011).

19. Lederman, S. J. & Klatzky, R. L. Hand movements: A window into haptic object recognition. Cogn. Psychol. (1987). doi:10.1016/0010-0285(87)90008-9

20. O’Doherty, J. E., Lebedev, M. A., Li, Z. & Nicolelis, M. A. L. Virtual active touch using randomly patterned intracortical microstimulation. IEEE Trans. Neural Syst. Rehabil. Eng. 20, 85–93 (2012).

21. Medina, L. E., Lebedev, M. A., O’Doherty, J. E. & Nicolelis, M. A. L. Stochastic Facilitation of Artificial Tactile Sensation in Primates. J. Neurosci. (2012). doi:10.1523/jneurosci.3115-12.2012

22. Ulrich, R. & Vorberg, D. Estimating the difference limen in 2AFC tasks: Pitfalls and improved estimators. Attention, Perception, Psychophys. (2009). doi:10.3758/APP.71.6.1219

23. Gescheider, G. A. Psychophysics: The fundamentals (3rd ed.). Psychophysics: The fundamentals (3rd ed.). (1997).

24. Stevens, S. S. On the psychophysical law. Psychol. Rev. (1957). doi:10.1037/h0046162

25. Crapse, T. B. & Sommer, M. A. Corollary discharge across the animal kingdom. Nature Reviews Neuroscience (2008). doi:10.1038/nrn2457

26. Ekman, Gös. Weber’s Law and Related Functions. J. Psychol. Interdiscip. Appl. (1959). doi:10.1080/00223980.1959.9916336

27. Dehaene, S. The neural basis of the Weber-Fechner law: A logarithmic mental number line. Trends in Cognitive Sciences (2003). doi:10.1016/S1364-6613(03)00055-X

28. Weber, E. H. E.H. Weber on the Tactile Senses. E.H. Weber on the Tactile Senses (2018). doi:10.4324/9781315782089

29. Kim, S. et al. Behavioral assessment of sensitivity to intracortical microstimulation of primate somatosensory cortex. Proc. Natl. Acad. Sci. 112, 15202–15207 (2015).

30. Bolanowski, S. J., Gescheider, G. A., Verrillo, R. T. & Checkosky, C. M. Four channels mediate the mechanical aspects of touch. J. Acoust. Soc. Am. (2005). doi:10.1121/1.397184

31. Lieber, J. D. & Bensmaia, S. J. High-dimensional representation of texture in somatosensory cortex of primates. Proc. Natl. Acad. Sci. (2019). doi:10.1073/pnas.1818501116

32. Dadarlat, M. C., Doherty, J. E. O. & Sabes, P. N. A learning-based approach to artificial sensory feedback leads to optimal integration. Nat. Neurosci. (2014). doi:10.1038/nn.3883

33. Thomson, E. E., Carra, R. & Nicolelis, M. A. L. Perceiving invisible light through a somatosensory cortical prosthesis. Nat. Commun. (2013). doi:10.1038/ncomms2497

34. Hartmann, K. et al. Embedding a Panoramic Representation of Infrared Light in the Adult Rat Somatosensory Cortex through a Sensory Neuroprosthesis. J. Neurosci. 36, 2406–2424 (2016).

35. Rognini, G. et al. Multisensory bionic limb to achieve prosthesis embodiment and reduce distorted phantom limb perceptions. J. Neurol. Neurosurg. Psychiatry (2018). doi:10.1136/jnnp-2018-318570

36. Shokur, S. et al. Expanding the primate body schema in sensorimotor cortex by virtual touches of an avatar. Proc. Natl. Acad. Sci. U. S. A. 110, 15121–15126 (2013).

37. Collins, K. L. et al. Ownership of an artificial limb induced by electrical brain stimulation. Proc. Natl. Acad. Sci. U. S. A. 114, 166–171 (2017).

38. D’Anna, E. et al. A closed-loop hand prosthesis with simultaneous intraneural tactile and position feedback. bioRxiv 8892, 262741 (2018).

39. Graczyk, E. L. et al. The neural basis of perceived intensity in natural and artificial touch. Sci. Transl. Med. 8, 1–11 (2016).

40. Fridman, G. Y., Blair, H. T., Blaisdell, A. P. & Judy, J. W. Perceived intensity of somatosensory cortical electrical stimulation. Exp. Brain Res. 203, 499–515 (2010).

41. Fetz, E. E. et al. Direct electrical stimulation of the somatosensory cortex in humans using electrocorticography electrodes: a qualitative and quantitative report. J. Neural Eng. 10, 036021 (2013).

42. Ifft, P. J., Shokur, S., Li, Z., Lebedev, M. A. & Nicolelis, M. A. L. A brain-machine interface enables bimanual arm movements in monkeys. Sci. Transl. Med. 5, (2013).

43. O’Doherty, J. E., Lebedev, M. A., Hanson, T. L., Fitzsimmons, N. A. & Nicolelis, M. A. L. A Brain-Machine Interface Instructed by Direct Intracortical Microstimulation. Front. Integr. Neurosci. 3, (2009).

44. Li, Z. et al. Unscented Kalman filter for brain-machine interfaces. PLoS One (2009). doi:10.1371/journal.pone.0006243

