## Supplementary material for "Creating a neuroprosthesis for active tactile exploration of textures": suuplementary material (from another version of this preprint: version 1, 2019-04-04)

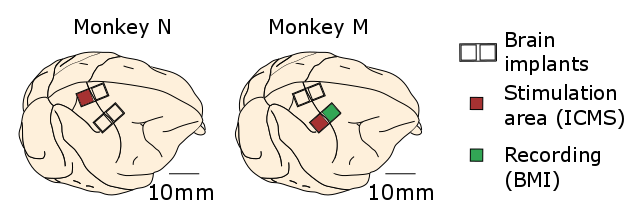

Fig S1. Electrode placements. Both monkeys were stimulated in their right primary somatosensory cortices: monkey M in arm region; monkey N in leg region. Recordings were made from the right primary motor cortex of monkey M for BMI control.

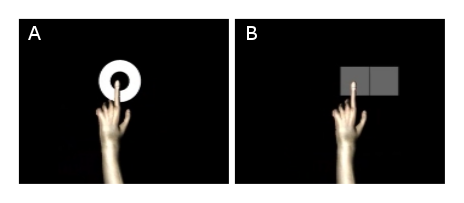

Fig S2. View of the monkey avatar arm projected on the monitor with the (A) central marker appearing at the onset of each trial, (B) two visually-identical targets appearing during the exploration task.

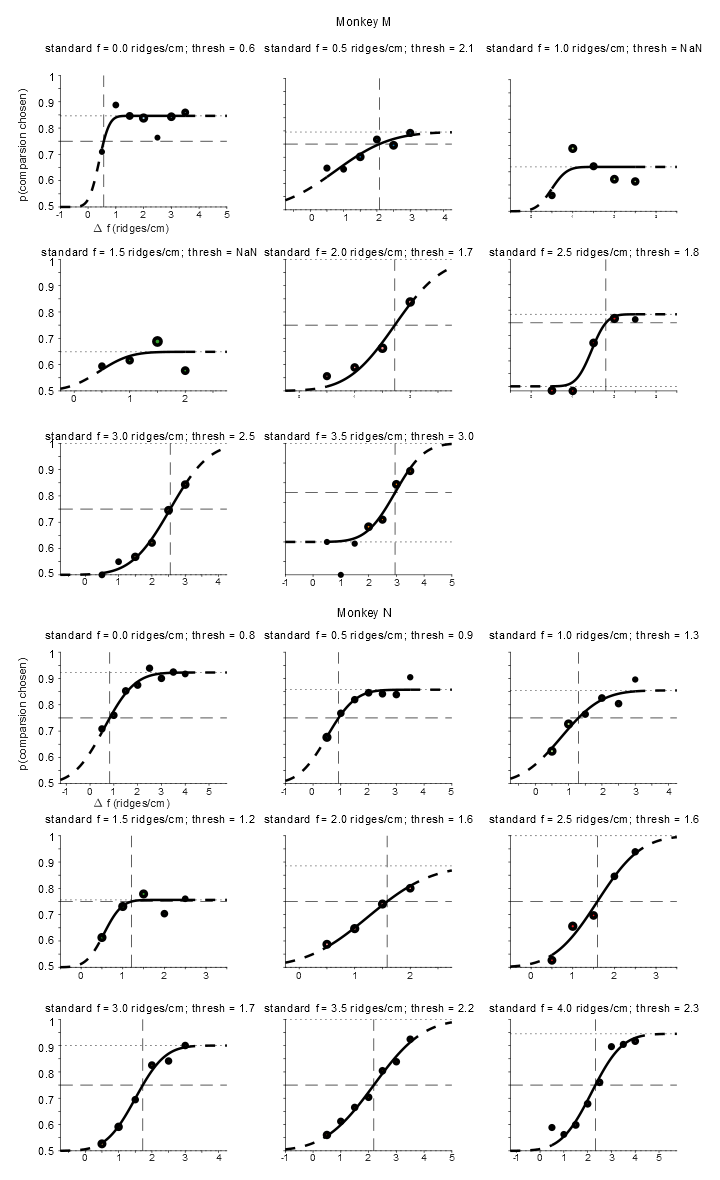

Fig S3. The psychometric functions used to calculate just noticeable differences (JNDs) for both monkeys and all target frequencies. Each graph shows, for a standard frequency, the probability of choosing a second comparison frequency as a function of the unsigned delta between the standard stimulus and the comparison stimulus. The line crossing the psychometric function at 75% probability (vertical dashed line) describes the JND. For monkey M in two cases (standard f = 1 and 1.5), the asymptotic performance was smaller than 75%, and the corresponding JND could therefore not be computed. Each filled circle indicates average performance at a tested comparison frequency; the size of the circle indicates the relative number of trials at that comparison frequency.

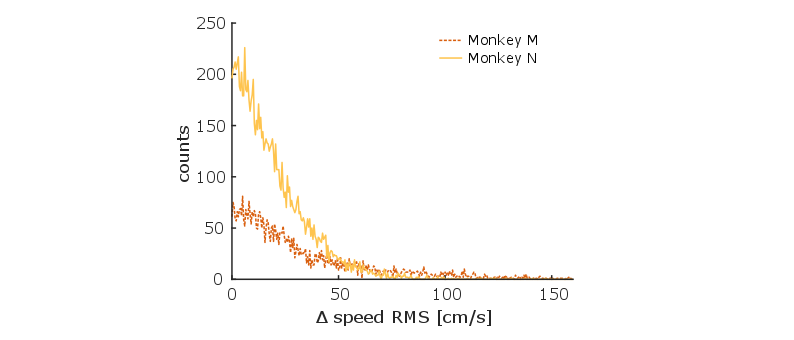

Fig S4. Distribution of the delta Root Mean Square (RMS) speed. For each trial (those with both correct and wrong outcomes) we calculated the difference of the RMS of the arm velocity when the monkey scanned each target (orange dashed line for monkey M and solid yellow for monkey N).

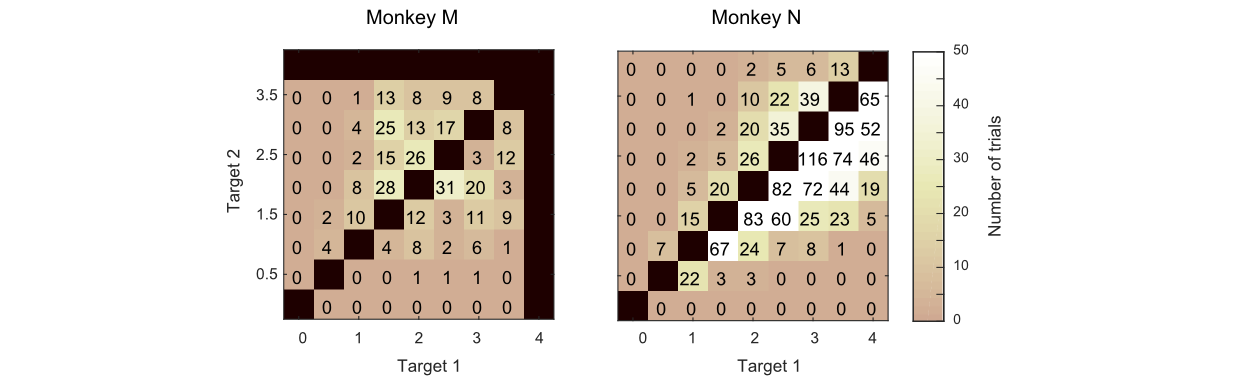

Fig S5. Number of apparent paradoxical trials factorized by frequency pair. A paradoxical trial indicates a trial where the target with highest delivered ICMS pulse rate was not the target with the highest spatial frequency. This can happen when the monkey’s scanning velocity is faster for the target with the lower spatial frequency.

Movie S1. Example of session for monkey N. Left side the video is a screen capture of the monkey’s monitor. The right side shows the experimenter’s monitor. The monkey did not see the visual cues for the different types of textures or the audio of the ICMS artifact. In this session the monkey controlled the avatar arm with a joystick. The monkey can scan both virtual objects and choose (by holding the avatar on top of one of the targets for 2 seconds) the target with the highest spatial frequency.

Movie S2. Example of session with monkey M. In this session the monkey controlled the avatar arm in BMI mode. The monkey can scan both virtual objects and choose (by holding the avatar on top of one of the targets for 1 second) the target with the highest spatial frequency. The rest of the experimental conditions were similar to the one described in movie S1.
